# Highly resolved genome assembly and comparative transcriptome profiling reveal genes related to developmental stages of tapeworm *Ligula intestinalis*

**DOI:** 10.1101/2023.07.25.550026

**Authors:** Masoud Nazarizadeh, Milena Nováková, Marie Drábková, Julian Catchen, Peter D. Olson, Jan Štefka

## Abstract

*Ligula intestinalis* (Cestoda: Diphyllobothriidae) is an emerging model organism for studies on parasite population biology and host-parasite interactions. However, a well resolved genome and catalogue of its gene content has not been previously developed. Here, we present the first genome assembly of *L. intestinalis*, based on Oxford Nanopore Technologies, Illumina and Omni-C sequencing methodologies. We use transcriptome profiling to compare plerocercoid larvae and adult worms and identify differentially expressed genes associated with these life stages. The genome assembly is 775.5 Mbp in size, with scaffold N50 value of 118 Mbp and encodes 27,256 predicted protein-coding sequences. Over 60% of the genome consists of repetitive sequences. Synteny analyses showed that the 10 largest scaffolds representing 75% of the genome display high correspondence to full chromosomes of cyclophyllidean tapeworms. Mapping RNA-seq data to the new reference genome we identified 3,922 differentially expressed genes in adults compared to plerocercoids. Gene Ontology analyses revealed overrepresented genes involved in reproductive development of the adult stage (e.g. sperm production) and significantly enriched DEGs associated with immune evasion of plerocercoids in their fish host. This study provides the first insights into the molecular biology of *L. intestinalis* and provides the most highly contiguous assembly to date of a diphyllobothriid tapeworm useful for population and comparative genomic investigations of parasitic flatworms.

## 1. Introduction

Parasites with multi-stage life cycles have evolved a number of mechanisms to facilitate transmission between the intermediate and definitive hosts and subsequent reproduction [1,2]. To understand parasite-host interactions on a molecular scale, it is necessary to develop genomic and transcriptomic resources that can elucidate the genetic underpinnings of the mechanisms associated with parasite adaptation. The genome of a parasite with a complex life cycle must contain genetic information that enables the parasite to survive and reproduce in vastly different environments with respect to metabolic requirements and strategies to evade the host immune system [3–8]. Therefore, comparative genomic analyses across several parasites can shed further light on the evolutionary adaptations [3,5,6,9,10]. Moreover, understanding the patterns of gene expression can provide key insights into the mechanisms underlying the regulation of physiological functions related to parasitic infection, within-host survival, and reproduction [9]. Therefore, an integrated approach combining both genomic and transcriptomic analyses promises to provide important insights into parasite biology and host-parasite interactions.

Significant strides in genomic research over the past decade have elucidated the genomic architecture of a broad spectrum of key species from the order Cyclophyllidea, predominantly the parasites of human or domestic animals [3,7,8,10–15]. Genomic and transcriptomic studies can reveal gene expression specific to each developmental stage in order to begin to understand the genes involved in parasitism and the features of parasitic life [5,7–9,16]. To date, genomic data exist for at least 17 cyclophyllidean tapeworms (NCBI; https://www.ncbi.nlm.nih.gov/ and WormBase ParaSite; [17]), five of which have highly contiguous, chromosome-level or fully complete assemblies [10–12,15,18]. In contrast, genomic resources for diphyllobothriidean tapeworms remain limited, with only four draft genome assemblies currently available [6,19,20]. Despite these disparities, both tapeworm orders share several life history characteristics, such as hermaphroditic adults parasitizing the intestines of vertebrates, and larval stages infecting invertebrate hosts. However, a stark ecological divergence is noticeable; whereas cyclophyllidean life cycles are typically bound to terrestrial ecosystems and hosts, the majority of diphyllobothriidean life cycles are associated with aquatic environments [21].

*Ligula intestinalis* is a common diphyllobothriidean cestode that has a nearly global distribution [22]. The parasite requires three hosts to complete its life cycle (electronic supplementary material, figure S1): a freshwater copepod as the first intermediate host, a wide array of fish species of the families Osmeriformes, Cypriniformes, Gobiiformes and Galaxiiformes as the second intermediate host, and a piscivorous bird as the definitive host [22–24]. The free-swimming coracidial larval stage of the parasite is consumed by planktonic copepods and develop into procercoid forms upon entering the haemocoel [23]. Once the copepod is consumed by fish, the procercoids mature into plerocercoid larvae in the abdominal cavity [22]. The parasites become sexually matured in the enteric tract of birds which consume the infected fish [22]. Compared to other tapeworms its adult stage is very short and only lasts for several days during which the eggs are dispersed by infected birds. Larval worms can be induced to develop sexually *in vitro* [22,25]. The tapeworm has been extensively used in for endocrinological, ecological, population genetic, phylogeographic and immunological studies [24,26–31].

The current study primarily aimed to provide the first highly contiguous genome assembly of *L. intestinalis*. Furthermore, we sought to investigate RNA transcription patterns to uncover the regulation of biological functions during the tapeworm’s life cycle, by establishing a comprehensive reference transcriptome sequence for both plerocercoid and adult stages of *L. intestinalis*. This research enriches our understanding of *L. intestinalis*’ molecular biology, offering a platform for further exploration in the field of population genomics for this species, as well as in comparative genomic studies of parasitic flatworms.

## 2. Materials and methods

### (a) Sampling and culturing

A total of 16 plerocercoids were collected from 15 roach (*Rutilus rutilus*) caught from river dams located in Želivka, Most and Klíčava in the Czech Republic according to previously described methods [31] (electronic supplementary material, table S1). To maintain uniform conditions and minimise variability in transcriptome expression, only 1-year old fish infected with a single tapeworm were selected from the Most and KlíČava water bodies (electronic supplementary material, figure S1). Whole genome sequencing was conducted on one sample obtained from a roach carrying two *L. intestinalis* parasites. The sample collected from the river dam in Želivka was rinsed with physiological saline solution (0.9% sodium chloride) and stored at -80 °C. Transcriptome sequencing was done using samples taken from anterior, central, and posterior parts of three living plerocercoids. Worms were removed from the fish in an aseptic environment, rinsed with Ultra Pure DNAse/RNase-free water (Ambion, Austin, Texas, USA) and cut into 5×5 mm square sections. Sections were immersed in RNAlater (Ambion, Austin, Texas, USA) at 4 °C overnight and stored at −80 °C prior to RNA extraction. Adult worms were grown in the lab using 12 plerocercoids transferred to the laboratory in physiological saline solution. Worms were paired and placed in six bottles containing 250 ml of culture medium prepared using 25 mM HEPES buffer (Sigma, Taufkirchen, Germany), Minimum Essential Medium with Earle’s salts, additives (1 ml/litre antibiotic ⁄antimycotic solution (Sigma), 6.5 g D-glucose) and L-glutamine. Adjustment of medium pH to 7.5 was done by adding NaOH. Culture bottles were incubated at 40 °C under constant agitation at 70 rpm. All six pairs of parasites were successfully cultured until they started producing eggs. Three adult parasites were randomly selected from three different pairs and samples were taken from their anterior, central, and posterior parts for RNA analysis as described for the plerocercoids.

### (b) DNA extraction and sequencing

DNeasy tissue kit (Qiagen, Germany) was used to retrieve high molecular weight (HMW) genomic DNA from plerocercoid larvae according to manufacturer instructions. The amount of extracted DNA was measured by a fluorometer (Qubit DNA BR Assay Kit, Invitrogen, USA) and DNA integrity assessed via gel electrophoresis. Two different strategies were taken to obtain data for genome assembly. First, we used Oxford Nanopore sequencing to obtain long DNA reads. The library was prepared and sequenced commercially (Roy J. Carver Biotechnology Center of the University of Illinois at Urbana-Champaign). HMW gDNA was converted into four Nanopore libraries with the 1D (SQK-LSK109) library kit and loaded onto a FLO-MIN106 R9.4.1 SpotON flow cell for 72 hs and sequenced in the GridIONx5 sequencer (Oxford Nanopore Technologies). The albacore algorithm (read_fast5_basecaller.py, ONT Albacore Sequencing Pipeline Software v2.3.1) was used for base calling all FAST5 files, resulting in 7,574,939 reads (average read length 5,699 bp, max. 123,333 bp, min. 5,402 bp) totalling 43 Gb of data (electronic supplementary material, table S2). To correct ONT reads another library using the same HMW sample as above was constructed for shotgun sequencing with the Hyper Library construction kit from Kapa Biosystems (Roche, Basel, Switzerland) and sequencing was performed using an Illumina NovaSeq at the Roy J. Carver Biotechnology Center of the University of Illinois at Urbana-Champaign. This sequencing yielded 485,493,102 paired-end 150 bp reads for a total of 146 Gbp (electronic supplementary material, table S2).

Second, to further improve genome contiguity we produced a chromosome conformation capture library [32]. An Omni-C proximity ligation library was prepared commercially (Dovetail Genomics, USA) using a flash frozen *L. intestinalis* sample. The library was sequenced using Illumina NovaSeq 150bp (106 Gbp, approximately 24X coverage) by the same commercial laboratory (electronic supplementary material, table S2).

### (c) RNA extraction and sequencing

We used RNA-seq to facilitate annotation of the coding regions of the genome and to investigate differences between plerocercoid and adult stages of the life cycle, as well as among three different regions of the worms. Samples were taken from the anterior, middle and posterior regions of three adult and three plerocercoid worms (18 samples in total). Total RNA extraction was done according to Chomczynski and Sacchi [33] by the acid guanidinium thiocyanate-phenol-chloroform method using Trizol® reagent (Invitrogen, Carlsbad, CA, USA). Integrity and yield were respectively evaluated in an Agilent Bioanalyzer 2100 (Agilent Technologies, USA) and a spectrophotometer (NanoDrop 1000; Thermo Scientific, USA). RNA samples were commercially processed into cDNA libraries and sequenced using Illumina Novaseq 150 PE read technology (Novogene UK and GeneWiz USA). The results were exported as FASTQ files.

### (d) Genome assembly

As a first step in the genome assembly, all FASTQ ONT files were merged into a single file and Flye v2.4 [34] was used to assemble ONT sequences. A total of 137 scaffolds were generated during the assembly process. The largest scaffold was 2,329,270 bp, the scaffold N50 was 320,313 bp and the mean coverage was 55x. The assembly was polished with Illumina 150 bp PE reads (60x coverage) using ntEdit [35]. Then, HiRise scaffolding (commercial service by Dovetail Genomics LLC) allowed for improvement of the scaffold contiguity using the OMNI-C library. Prior to annotation, we excluded scaffolds with coverage < 10× and length < 500 bp that constituted less than 1% of our data. Moreover, their exclusion did not omit any contigs containing hits with the transcripts from *L. intestinalis*. We used Phyloligo version 1.0 [36] to check for contamination in the genome assembly. To evaluate the quality of the genome assembly, we utilized GenomeQC [37] and QUAST v4 [38]. Genome completeness was estimated via BUSCO v5 (Benchmarking Universal Single-Copy Orthologs), as well as searching for BUSCO groups in the Metazoan database [39]. The whole genome sequencing data from our study is accessible on GenBank, associated with the Bioproject number (available upon acceptance of the paper).

### (e) Prediction of coding genes

Repeat families were first identified and classified using RepeatModeler v2.0.1 [40], a package that utilises RepeatScout v1.0.6 and RECON v1.08 to detect repeated sequences in genomes. The output from RepeatModeler was then processed using RepeatMasker v 4.1.0, allowing the repeated sequences to be identified, classified, and masked in the assembly file. Upon completion of the repeat masking, an initial ab initio gene prediction model for *L. intestinalis* was generated, using coding sequences from *Dibothriocephalus latus, Schistocephalus solidus, Sparganum proliferum, Spirometra erinaceieuropaei*, and *Taenia solium* as references. This gene prediction model was optimised through six rounds of prediction adjustments, made in AUGUSTUS v2.5.5 [41]. Concurrently, a separate ab initio gene prediction model was trained using the same set of coding sequences in SNAP v2006-07-28 [42].

To further improve the accuracy of the gene predictions, RNA sequences were aligned to the assembled genome using the STAR aligner software v2.7 [43]. Intron hints were also obtained using the bam2hints tool from AUGUSTUS, which were used to refine the RNA sequence alignment to our genome. Gene prediction was then performed in this repeat-masked reference genome, using a combination of AUGUSTUS, SNAP, and MAKER. These programs used the intron-exon boundary hints from the RNA sequences to guide the prediction process. Swiss-Prot peptide sequences from the UniProt database and from *D. latus, S. solidus, S. proliferum, S. erinaceieuropaei*, and *T. solium* were incorporated into the MAKER pipeline to inform peptide generation.

Genes that were not predicted by both AUGUSTUS and SNAP were excluded from the final gene set. The MAKER pipeline assigned Annotation Edit Distance (AED) scores to the predicted genes as a measure of prediction performance. Subsequently, BLAST searches were conducted in the UniProt database to gather further information about the function of our predicted genes. In parallel, tRNA prediction was carried out using tRNAscan-SE v 2.05 [44]. Lastly, TransposonPSI [45] was used to identify potential transposons within the predicted protein set and to discover regions of transposon homology within the genome sequence.

### (f) Functional annotation of the gene models

We carried out the functional annotation of the predicted genes using the OmicsBox v2.2.4 software [46], through the BLASTx function [47]. We set an E-value cut-off of 10^-5 and cross-referenced with multiple databases including eggNOG (Orthologous groups of genes; [48], InterProScan [49], Swiss-Prot [50], and the NCBI non-redundant protein sequences database (Nr; [51]). Furthermore, to enhance the quality of our gene annotations, we individually compared the protein sequences using the Pfam database [52], setting a HMMER E-value cut-off at 10^-10.

### (g) Mitochondrial genome

Reconstruction of the mitochondrial genome was done using Illumina reads in MITObim v.1.67 [53]. *L. intestinalis* (NC039445) mitochondrial genes were used as queries by BLASTX to identify mitochondrial fragments in the assembly. To extend the fragments, we mapped the Illumina short reads to the assembly in MITObim. The MITOS web server was used to annotate protein-coding, ribosomal RNA and tRNA genes. Artemis genome annotation tool [54] was used to manually curate annotations and assemblies based on sequence similarity to other published mitogenomes.

### (h) Orthologous genes and species and gene family tree

Orthologous and duplicate genes were identified via phylogenetic analysis using OrthoFinder v2.5.2 (Emms and Kelly, 2015, 2019). Then, the proteomes predicted in the *L. intestinalis* genome were compared to those in other species of Diphyllobothriidea (*S. solidus* (NCBI, BioProject: PRJNA576252), *S. erinaceieuropaei* (NCBI, BioProject: PRJEB35375), *D. latus* (NCBI, BioProject: PRJEB1206)) and Cyclophyllidea (*Taenia multiceps* (NCBI, BioProject: PRJNA307624), *T. saginata* (NCBI, BioProject: PRJNA71493), *Echinococcus granulosus* (NCBI, BioProject: PRJNA754835), *Hymenolepis microstoma* (NCBI, BioProject: PRJEB124), *H. nana* (NCBI, BioProject: PRJEB508) and *H. diminuta* (NCBI, BioProject: PRJEB30942)). For increased accuracy, DIAMOND blast (E-value □ < □ 1e−5) in OrthoFinder was used [55,56]. The Markov Cluster algorithm (MCL) was used for sequence similarity and clustering. MAFFT v.7 [57] was utilised for multiple protein sequence alignment, and FastTree2 v2.1 [58] for inference of maximum likelihood gene trees for each orthogroup. OrthoFinder used a concatenated alignment of single copy orthogroups to infer a species tree with at most one gene per species. The tree was reconstructed using the Species Tree Root Inference from Duplication Events (STRIDE) algorithm [59]. The results were plotted using the cogeqc package in R [60,61].

### (i) Synteny analyses

We compared synteny between the *Ligula* genome and two chromosome-level tapeworm assemblies available: *H. microstoma* (BioProject: PRJEB124) and *E. granulosus* (BioProject: PRJNA754835). Due to the fragmented state of the other Diphyllobothriidea genomes, synteny analysis was performed with only the two most contiguous assemblies of *S. erinaceieuropaei*, and *S. proliferum* (table 1) using SyMAP v5.1.0 (Synteny Mapping and Analysis Program) [62,63]. For the synteny analysis, we omitted all scaffolds shorter than 1 Mbp and then aligned sequences using MUMmer [64]. The raw anchors generated by MUMmer were pooled into (putative) gene anchors, followed by filtering with a reciprocal top-2 filter (top_n = 2).

**Table 1.**
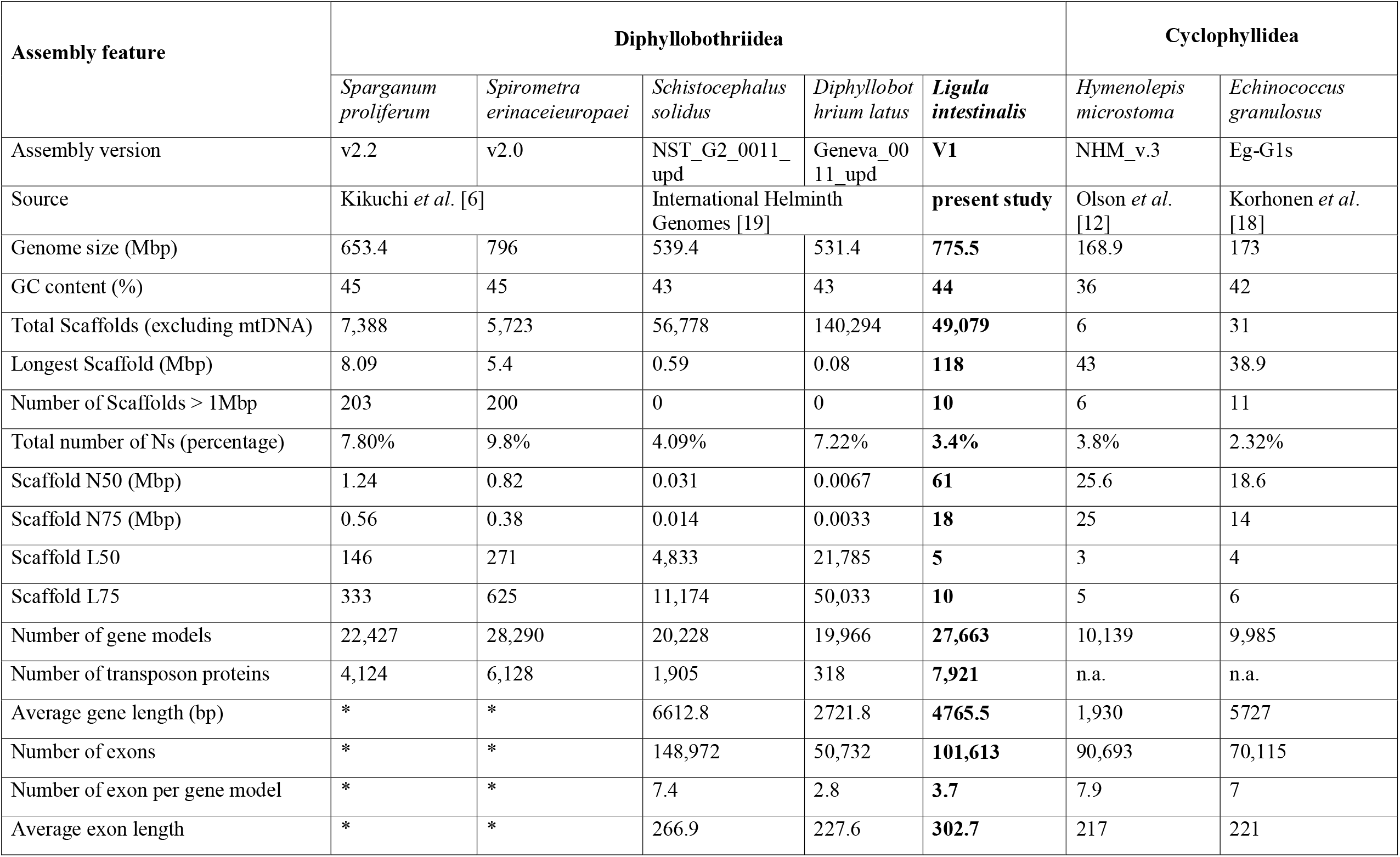
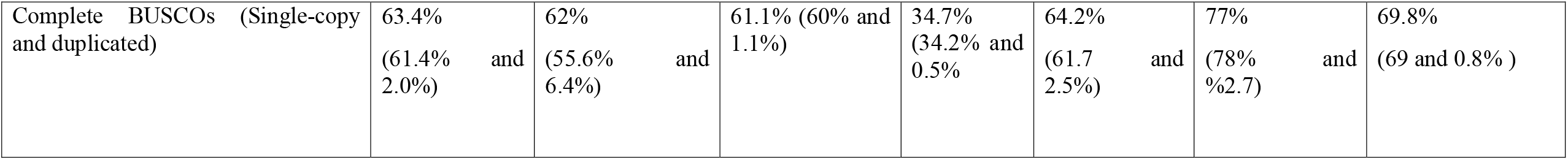
Genome assembly metrics of *L. intestinalis* and four other tapeworms in the order Diphyllobothriidea, along with two representative complete genomes from the order Cyclophyllidea. n.a. - data not analysed *Data mis-reported in Kikuchi *et al*. [6] due to missing intron information.

### (j) Differentially expressed genes (DEGs)

A total of 18 RNA-seq samples were used to compare transcriptomic differences between plerocercoid and adult worms, and among the anterior, middle and posterior regions of both life stages (three replicates for each body part and life stage). Low quality reads were initially discarded from all samples using Trimmomatic v0.33 [65]. Following trimming, paired-end reads (electronic supplementary material, table S1) were mapped to the genome of *L. intestinalis* using STAR. We also employed Samtools v1.10 [66] to eliminate reads that mapped to multiple loci. The filtered SAM files were sorted and converted to BAM format and the mapped reads counted for genomic features using featureCounts v1.06 in the Subread package [67]. Electronic supplementary material, table S3 provides mapped reads per sample replicate and gene model. The Trimmed Mean of M values (TMM) method was applied to normalise the raw counts. We used DiCoExpress R script [68] to estimate differential expression of genes between two life stages (plerocercoid and adult) and categorised them based on the contrasts between the different regions of the body. The R script uses generalised linear models in the package edgeR [69]. To identify DEGs, we used criteria of p-value < 0.05 and absolute log-fold change > 1. In order to evaluate the general similarities and differences among all the transcriptomes, plotMDS function in the edgeR was utilised to plot the first two principal components.

### (k) Gene Ontology and enrichment analysis

The clusterprofiler package [70] in R was utilised to perform a hypergeometric test for functional annotation of Gene Ontology (GO). The Benjamini-Hochberg false discovery rate adjustment of the p value was applied before comparisons were made in the enrichment analysis with a threshold value of 0.05. Dot plots and enrichment maps for over-and under-represented genes were then generated using the DOSE R package [71] to visualise the results. The outcome of the enrichment analysis was juxtaposed with the findings from the DicoExpress analysis. GOs were summarised using REVIGO for sets of DEGs with a threshold of 0.4 and SimRel as a measure of similarity [72].

## 3. Results

### (a) Genome metrics

We sequenced genomic DNA and first generated a nanopore-based 778 Mbp assembly, which was then improved with Omni-C data (electronic supplementary material, table S2). Using this combination of sequencing technologies, a 775.5 Mbp draft genome was assembled, representing one of the largest genome estimates yet known for diphyllobothriidea tapeworms, which range from 531-796 Mbp. Table 1 compares the assembly and genome metrics of *Ligula* with four other genomes in the same tapeworm order (*S. proliferum, S. erinaceieuropaei, S. solidus*, and *D. latus*) and two representative chromosome assemblies of Cyclophyllidea (*H. microstoma* and *E. granulosus*). The new *Ligula* genome had the best results to date for scaffold N50 (61 Mbp) and L50 (5) in comparison to the other four species, respectively. The GC content of the draft assembly is 44%, which is similar to those species (43%-45%). Out of 954 BUSCOs, 613 (64.2%) were complete, 89 (9.3%) were fragmented, 24 (2.5%) complete and duplicated, and 252 (26.5%) were missing from the assembled genome. 61.1% of the genome was found to comprise repetitive sequences, including 7,921 unique transposon protein sequences (table 1 and electronic supplementary material, table S3).

### (b) Orthologous genes and gene duplications

A total of 198,304 proteins from 11 cestode species were placed into 165,590 (83.15%) gene families in orthogroups and 32,714 (16.5%) unassigned singletons. Figure 3 depicts a species tree for five diphyllobothriidean and six cyclophyllidean species and demonstrates gene duplication events for each node and tree branch. *L. intestinalis* separated from *D. latus* with high support value and 767 gene duplication events. The *L. intestinalis* proteome (27,256 proteins) was encoded by 19,104 (70.8%) gene families (figure 1A), 1,829 of which were shared across all diphyllobothriidean and cyclophyllidean species (figure 1B). Among the proteins, 6,677 (24.5%) were specific to the species or singletons. Also, the highest number of species-specific duplication and species-specific orthogroups were observed in the *Ligula* genome (figure 2A). Lastly, the genome had the highest and the lowest number of orthologos compared to *D. latus* and *T. multiceps*, respectively.

**Figure 1.**
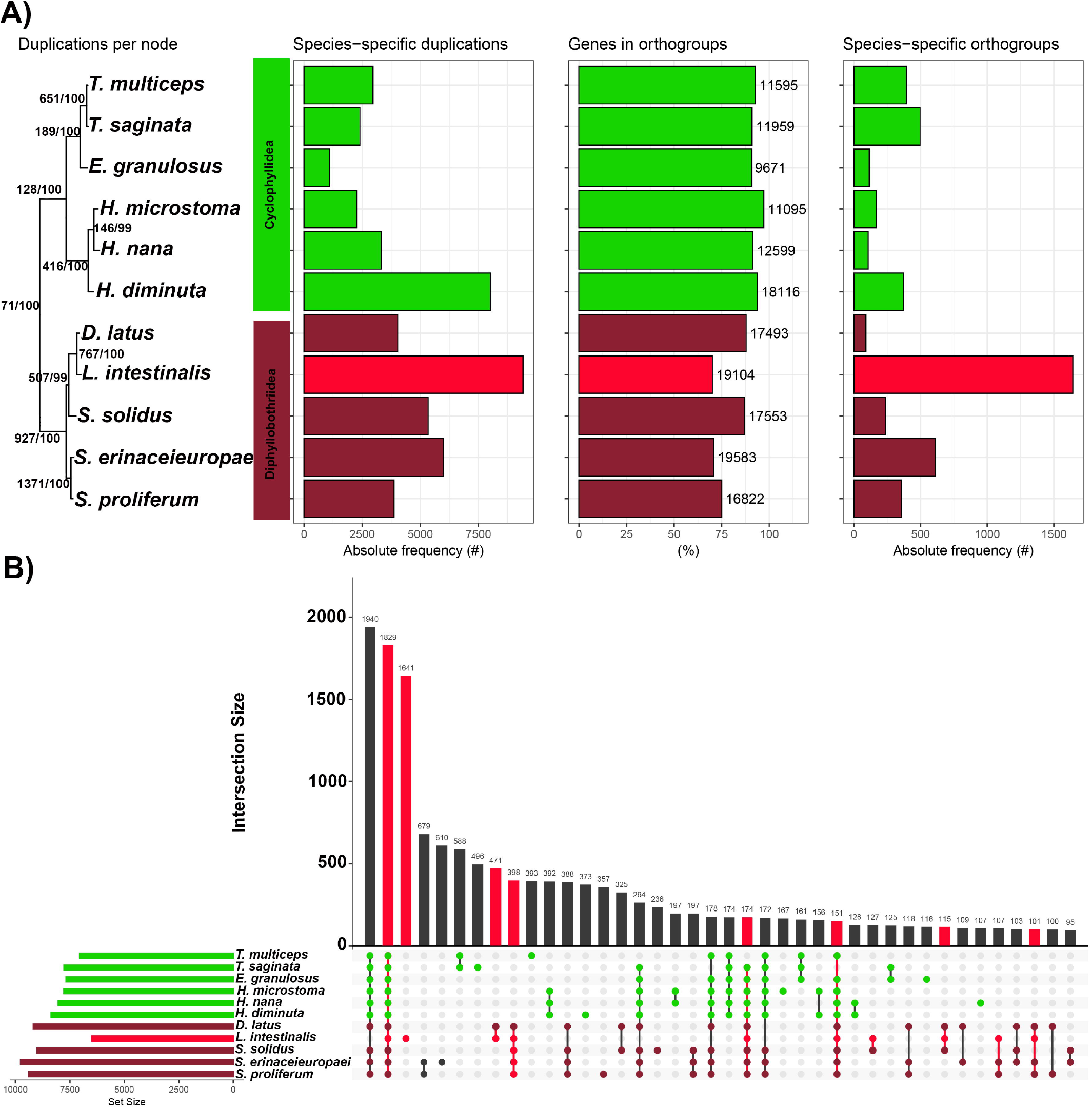
Gene duplication and orthology in diphyllobothriidean and cyclophyllidean cestodes. A) Summary of OrthoFinder analysis on five available genomes of diphyllobothriidean species and six representative genomes of cyclophyllidean tapeworms. The species tree was reconstructed using a concatenated alignment of 226 single copy orthogroups, with the number of top branches corresponding to gene duplication events and bootstrap value. The horizontal bar plots show gene duplication events per species, percentage of genes in orthogroups, and the number of species-specific orthogroups. The tree is rooted with a cyclophyllidean tapeworm *E. granulosus* B) UpSet plot of the intersection of gene families in diphyllobothriidean and cyclophyllidean tapeworms.

**Figure 2.**
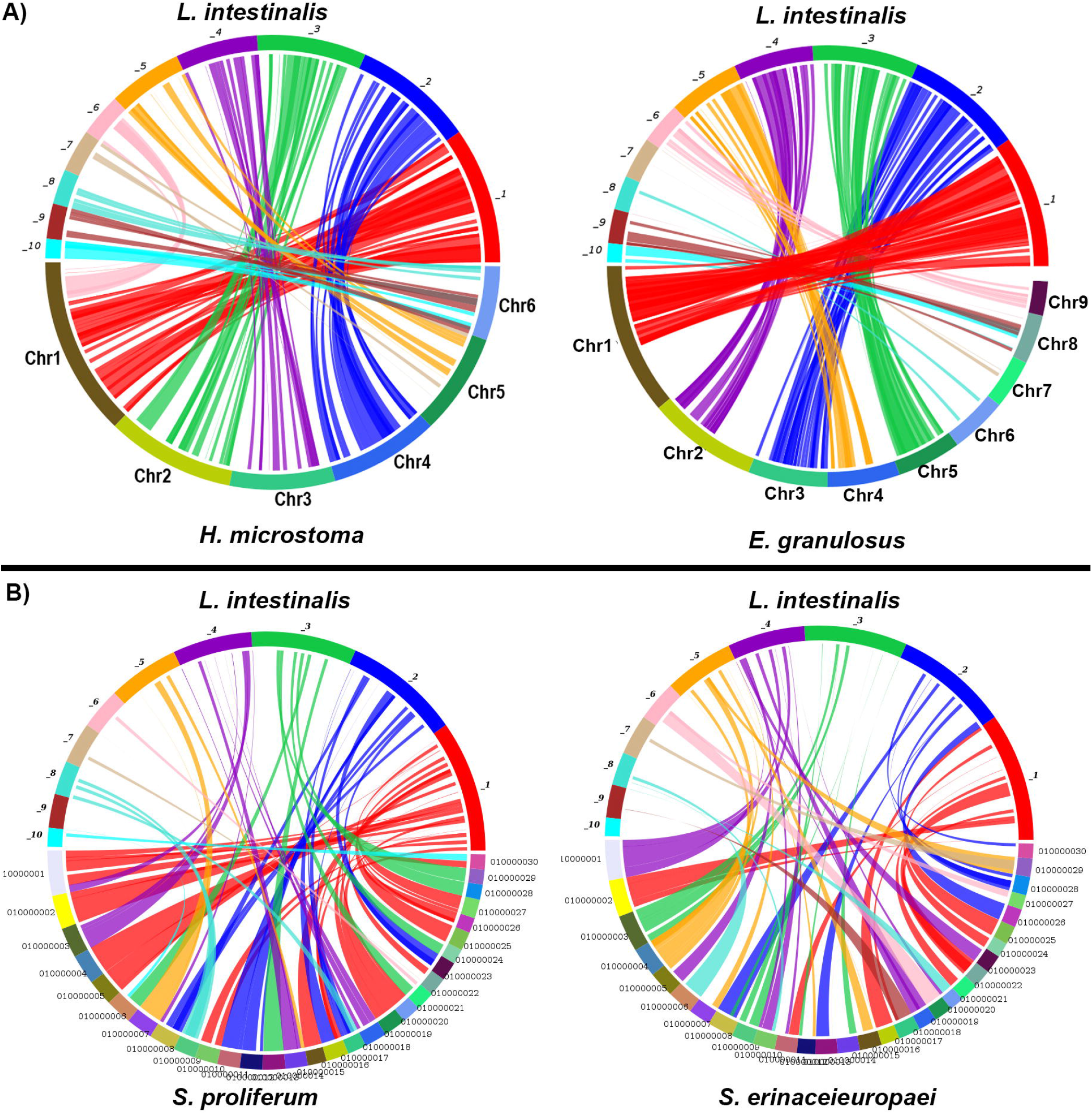
Syntenies of *L. intestinalis* and cestodes of the cyclophyllidean and diphyllobothriidean orders. A) Chromosomal synteny between *L. intestinalis* and two representative complete genomes in Cyclophyllidea (*H. microstoma* and *E. granulosus*) is highly conserved. B) Synteny plots between *L. intestinalis* and *S. proliferum* and *S. erinaceieuropaei* genomes demonstrate higher contiguity of *L. intestinalis* genome.

**Figure 3.**
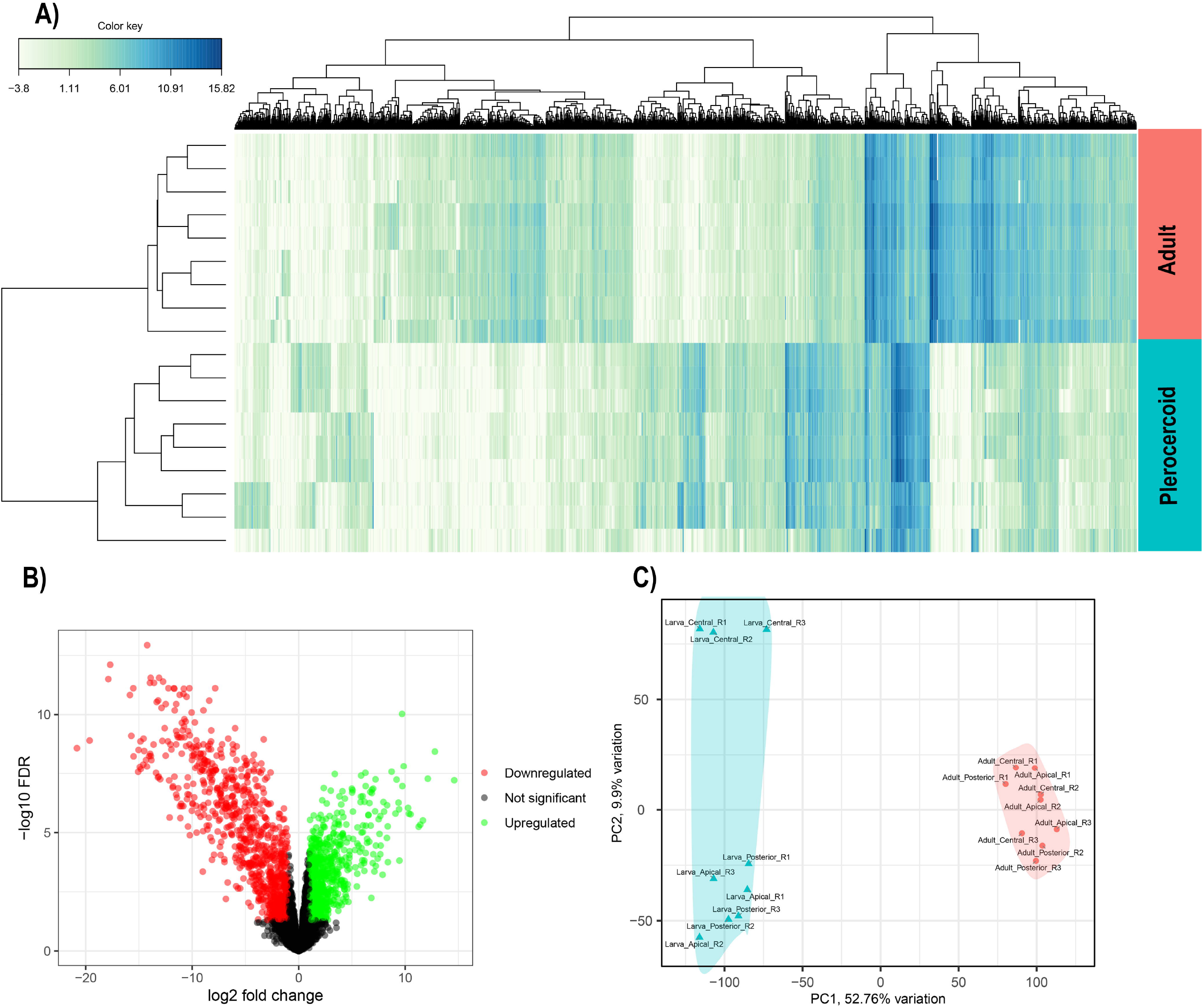
Differential gene expression in *L. intestinalis*. A) Heatmap of the differentially expressed (DE) genes between plerocercoids and adults. B) Volcano plot of up/down-regulated and non-DE transcripts between plerocercoids and adults. Red and green dots correspond to genes with significantly differentially expressed transcripts, while transcripts whose expression levels were not significant are depicted as blue dots. C) Principal component analysis for clustering of sample replicates with relative variances across PC1 and PC2. Larval and adult samples are colour coded as in plot A.

### (c) Comparative Synteny Analysis between *Ligula* and Other Genomes

We analysed chromosome-level synteny among the ten longest scaffolds of the *Ligula* assembly with the six chromosomes of *H. microstoma* and nine chromosomes of *E. granulosus*. Extensive preservation of synteny is evident when the first six scaffolds are compared with the chromosome assemblies of the two species. Large regions of these scaffolds are aligned with single chromosomes in both species. However, scaffolds eight to ten were predominantly mapped to chromosome six in *H. microstoma* and to chromosomes eight and nine in *E. granulosus* (figure 2). We also mapped the first 30 scaffolds of *S. erinaceieuropaei* and *S. proliferum* to the 10 scaffolds of *L. intestinalis* (figure 2B). The 10 longest scaffolds cover 75% of the *Ligula* genome, whereas in *Sparaganum* and *Spirometra* only 60% and 45% genes were covered by their 10 longest scaffolds, respectively. We also compared the 10 longest scaffolds of the *Ligula* genome with 200 and 203 scaffolds of *S. erinaceieuropaei* and *S proliferum*, respectively (electronic supplementary material, figure S2, all scaffolds longer than 1Mbp). Analysis of pairwise collinearity of orthologous blocks on the scaffolds showed a stronger similarity between *L. intestinalis* and *S. proliferum* (342 blocks and 16089 anchors) than between *L. intestinalis* and *S. erinaceieuropaei* (240 blocks and 10,657 anchors). The longest scaffold (118 Mbp) in the *L. intestinalis* genome corresponds to 41 and 42 scaffolds in *S. erinaceieuropaei* and *S. proliferum*, respectively.

### (d) Differential Gene Expression and Gene Ontology Patterns

Our transcriptome analyses identified 3,922 differentially expressed genes between the two life stages (electronic supplementary material, figure S3, figure 3A and electronic supplementary material, table S4), comprising 2,301 up-regulated and 1,621 down-regulated genes in the plerocercoids. Interestingly, RNA expression patterns revealed that larvae and adults clearly diverged from each other by the first dimension of PCA (52.76% variance explained), and the central part of larval worms separated from its posterior and anterior parts by the second dimension of PCA (9.9%) (figure 3C). Moreover, in analysis of the relationship between different parts of the body and life stage, the central and posterior parts of larvae and adults demonstrated 1,856 DEGs (1,270 up-regulated and 586 down-regulated) and 1,573 DEGs (945 up-regulated and 628 down-regulated), respectively (electronic supplementary material, figure S2 and table S4).

Up- and down-regulated DEGs were analysed using the GO enrichment method to identify the characteristics of the DEGs between adults and plerocercoids. Results using both DiCoExpress and clusterprofiler were similar, however, DiCoExpress not only encompassed all the enriched genes identified by clusterprofiler but also provided additional insights by revealing an increased quantity of overrepresented and underrepresented genes (electronic supplementary material, table S5 and table S6).

The enrichment analysis from clusterprofiler revealed that the biological process of the 275 overrepresented DEGs were mainly enriched in the cytoskeleton organisation (94 DEGs), cilium organisation (82 DEGs), microtubule-based movement (70 DEGs), microtubule-based process (12 DEGs), axoneme assembly (9 DEGs), and organelle assembly (8 DEGs) (figure 4). Moreover, the cellular components of the upregulated genes revealed 262 DEGs corresponding to cilium (104 DEGs), cytoskeleton (79 DEGs), cytoplasm (55 DEGs), microtubule cytoskeleton (16 DEGs), and cytoplasmic region (8 DEGs). In addition, molecular function was associated with 123 upregulated DEGs, which mainly included protein binding (62 DEGs), cytoskeletal motor activity (26 DEGs), structural constituent of cytoskeleton (11 DEGs), calcium ion binding (12 DEGs), and GTP binding (12 DEGs) (figure 4). The 59 downregulated DEGs in biological processes were mainly involved in monoatomic cation transmembrane transport (8 DEGs). The corresponding cellular component of downregulated DEGs included extracellular space (28 DEGs), caveola (7 DEGs), membrane (54 DEGs), and cellular anatomical entity (16 DEGs). Finally, the molecular function of downregulated DEGs revealed overrepresented genes in actin binding, transmembrane signaling receptor activity, serine-type endopeptidase activity, protein binding, monoatomic ion transmembrane transporter activity, inorganic anion transmembrane transporter activity, and gated channel activity (figure 4).

**Figure 4.**
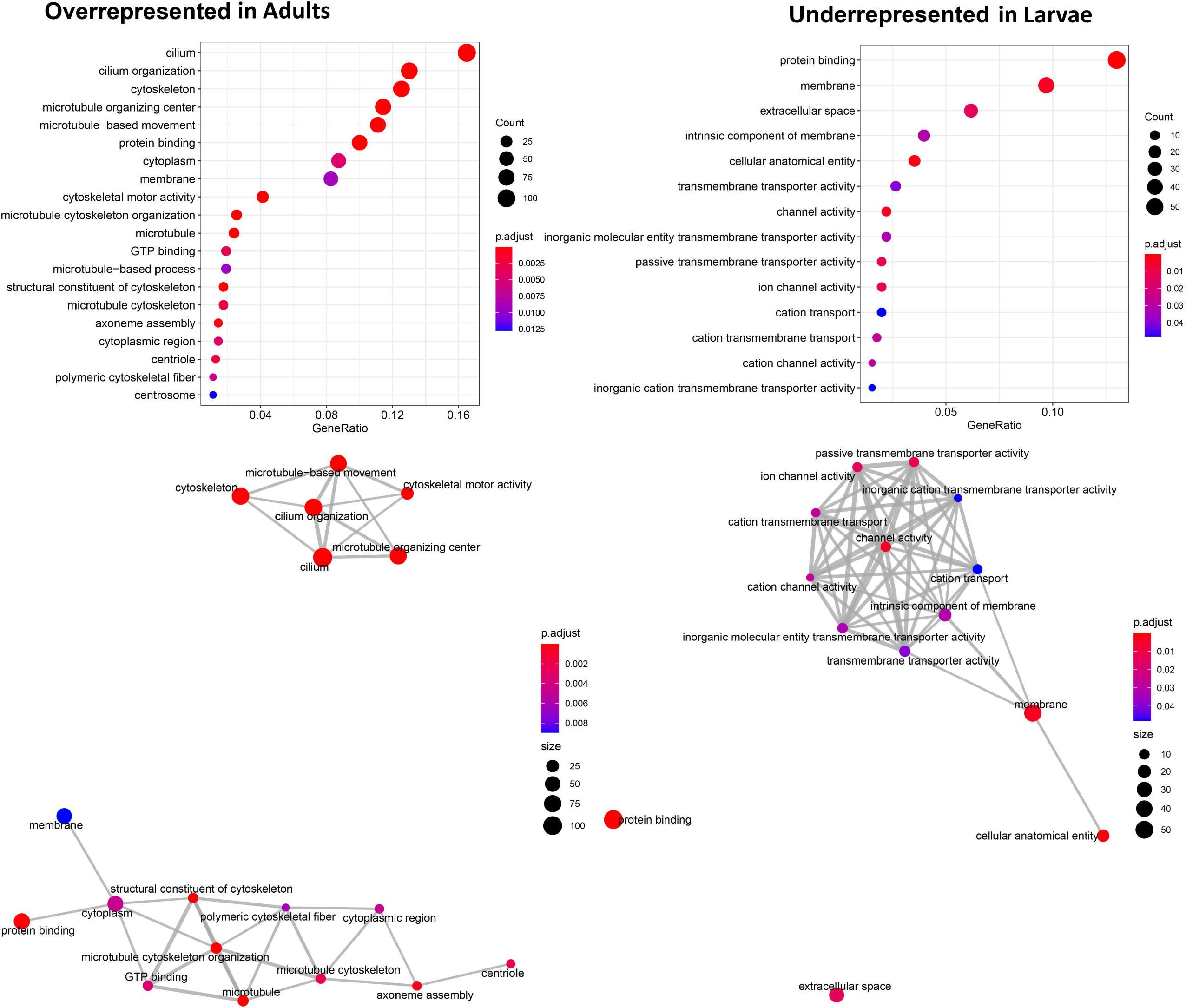
GO enrichment analysis of DEGs between larva and adult. Dot plot and directed acyclic graph show the up-regulated and down-regulated genes GO terms (FDR <0.05) of biological processes, molecular functions, and cellular components to be enriched between larvae and adults.

## 4. Discussion

Hybrid assembly facilitates more accurate and contiguous genome assemblies, which allow for improved estimates of the genome content and the ability to analyse syntenic relationships and other features of genome architecture [12]. Recent genomic studies on cestodes have used multiple sequencing technology platforms to improve previously published genomes (e. g., 7,13,14,16,22,74). In this study, the *L. intestinalis* genome was constructed using a hybrid assembly, and comparative genomics was conducted using all other available genomes representing the Diphyllobothriidea and six genomes representing the Cyclophyllidea. This study revealed substantial conservation of synteny between the first six scaffolds of the *Ligula* genome and chromosome level assemblies of *H. microstoma* and *E. granulosus*. In addition, the ten longest scaffolds constitute 75% of the total genome assemblies and 66% of the total number of genes. Assembly quality metrics (NX and LX) indicate high contiguity for the new *Ligula* genome in comparison to other diphyllobothriidean genomes. Consequently, the *L. intestinalis* genome acquired in this study allows for a more reliable estimate of the genome’s contents and size for this tapeworm.

### (a) Genome size, repetitive content and gene duplications

In the newly assembled genome, we discovered a substantial proportion of repetitive sequences (electronic supplementary material, table S3; 61.1%), which aligns with previous findings of high repetitive sequence content in three other diphyllobothriidean species (*D. latus, S. proliferum*, and *S. erinaceieuropaei*). In addition, sequenced diphyllobothriidean species consistently have a larger size and number of species-specific genes compared to sequenced cyclophyllidean tapeworms [6,12]. In addition to the increased repeat content inflating the genome size in Diphyllobothriidea, our study and the previous studies have shown that diphyllobothriideans have experienced more expansions or fewer contractions of gene families compared to cyclophyllidean tapeworms [6,13]. These factors could explain the observed differences in genome size between these two major tapeworm groups and suggest that cyclophyllidean genomes have become streamlined.

Reconstruction of the phylogenetic relationships among the five diphyllobothriidean species recovered a generally well-supported tree that is congruent with a previous genomic study not including *L. intestinalis* [6]. The finding of *L. intestinalis* and *D. latus* being recovered as sister taxa to each other and in turn sister clades to *S. solidus* is in line with a study based on more samples but fewer genetic loci [74]. In accordance with the phylogeny, the highest number of genes in orthologs was found between *L. intestinalis, D. latus*, and *S. solidus*. Higher numbers of orthologous genes indicate a more recent common ancestor and a shorter divergence time, whereas lower numbers indicate a more ancient common ancestor and a longer divergence time [75–77]. The results indicated a relatively high frequency of gene duplication during the divergence of *L. intestinalis* and *D. latus*, as well as between *S. proliferum* and *S. erinaceieuropaei*. Such frequent gene duplication events possibly played a major role in the speciation processes [78]. Additionally, *L. intestinalis* exhibits a high number of species-specific duplications (figure 1). Although the implication of these duplications remains to be tested, it could be suggested that it allows *L. intestinalis* a greater potential for functional diversification and host specialization [79–81] compared to the other four species.

### (b) Transcriptomics and stage-specific gene expression

We conducted the first comprehensive analysis of the plerocercoid and adult stages of *L. intestinalis* using RNA sequencing methods. Results show that more than 80% of all transcriptome reads from both life stages could be assigned to the new reference genome. This indicates that the RNA-seq data obtained are of high quality and can accurately quantify the expression level of genes [82]. Moreover, the results revealed different transcriptome patterns between the two life stages and identified numerous potentially DEGs associated with various biological processes, cellular components, and molecular functions. Interestingly, enrichment results showed that although *L. intestinalis* has the shortest adult life stage (3-7 days) among pseudophyllidean tapeworms, enabled by a plerocercoid stage that develops to the size of an adult inside the fish intermediate host [22,23]. It then undergoes significant physical and biochemical changes during the transition from the larval to adult phase, similar to other tapeworm species (e.g., 6,7,10,17,84). These changes include modulation of gene expression associated with various biological tasks such as reproduction, development and host survival.

The results of our enrichment analysis revealed an abundance of overrepresented genes associated with cilia and microtubules in adult tapeworms (electronic supplementary material, table S5, table S6, and figure 4). In agreement with the present study, Preza *et al*. [16] showed significantly overrepresented cilia and flagellum genes in a rodent tapeworm (*Hymenolepis microstoma*). These structures are essential for sperm production. Cilia, hair-like structures, facilitate sperm movement within the reproductive system [84], while microtubules provide structural support and contribute to the formation and organisation of the sperm tail or axoneme [85]. Correspondingly, our results also revealed several significantly enriched genes directly related to adult tapeworm sperm production (e.g. axoneme assembly, sperm motility, microtubule-based movement, and microtubule-based processes) (figure 4 and electronic supplementary material, table S5). Spermatogenesis is characterised by the formation of a flagellum that is integrated into the cytoplasmic extension of the spermatid, eventually leading to a mature spermatozoon. The spermatozoon has unique features, such as a single spiral anterior body, a cortical microtubule, and two axonemes with different lengths (the 9+”1” and 9+9+2 patterns), which distinguish the mature spermatozoon of *L. intestinalis* from other tapeworm species [86,87]. Therefore, studies of the candidate genes involved in axoneme assembly and microtubule function in *L. intestinalis* may provide valuable insights for evolutionary developmental relationships among tapeworms.

In addition, we discovered a prominent pattern characterised by a significant enrichment of the cytoplasmic region (electronic supplementary material, figure S2 and figure 5; cytoplasm, cytoskeleton, Golgi cisternae membrane, dynein complex) as a cellular component during the development of adult tapeworms. It is known that cytoplasm plays a crucial role in reproduction, nutrient uptake, and energy production in tapeworms [88–91]. Consistent with our findings, high levels of expressed transcripts have been reported for cytoskeletal proteins in multiple cestode species, particularly in diphyllobothriidean cestodes [8,10,92,93]. The cytoplasm of vitellocytes houses organelles such as ribosomes, the rough endoplasmic reticulum, and the Golgi apparatus, which are involved in the synthesis, modification, and packaging of yolk proteins. In accordance with that, Yoneva *et al*. [91] demonstrated that vitellocytes undergo significant cytoplasmic changes during vitellogenesis in *L. intestinalis* as they produce yolk components.

The present study also identified several over-represented genes associated with osmoregulatory balance in adult *L. intestinalis* (electronic supplementary material, table S5-S6). Osmoregulation is critical for many parasitic flatworms that live in their host’s intestine because they lack a digestive system and must therefore absorb nutrients from the intestinal contents of the host [94]. However, *L. intestinalis* uses the host intestine mainly as a means of sexual maturation and egg dispersal, while it is not thought to absorb nutrients during its short-lived adult stage [22]. On the other hand, possible differences in osmotic pressure between the intermediate host environment (plerocercoids in fish body cavity) and the bird intestine could explain the transcriptional differences between plerocercoids and adults even without the need for nutritional intake of the adult. Lastly, we should note that the *in vitro* conditions used to raise adults in this study may have resulted in an unnatural osmotic stress in the tapeworm. Consequently, further research is needed to conclusively determine the osmotic balance of *Ligula* tapeworms, with an emphasis on understanding the molecular and cellular mechanisms involved in osmoregulation.

Recent transcriptome analyses of several tapeworm species have identified an increased number of upregulated genes related to immune evasion in the larval stage compared to the adult [6–8,93]. Similarly, we identified DEGs significantly enriched in immune evasion of the plerocercoid larvae (electronic supplementary material, table S6). Consequently, combating the host immune system is critical for parasite’s survival and establishment in the fish intermediate host. Phylogeographic studies have revealed several evolutionary lineages of *L. intestinalis*, with each lineage exhibiting a high degree of host specificity for particular fish species groups [24,29,30]. Although the precise immune response of *L. intestinalis* is still unclear, strategies to evade the immune system may play a crucial role in the host specificity of different *Ligula* lineages and thus may evolve via directional selection. Focusing on sequential differences between evolutionary *Ligula* lineages in the DEGs identified here could help understanding the process of host-specialisation assumed to drive speciation in *Ligula* [24,29,30].

In conclusion, the high-quality genome and transcriptomic profiling of the larvae and adults provided by this study can serve as a starting point for research targeted at developmental biology, host-parasite co-evolution and immunogenetics in *L. intestinalis* and other diphyllobothriidean tapeworms. Although the current assembly with its 10 longest scaffolds covering 75% of the genome length represents the most contiguous genome available for any non-cyclophillidean tapeworm, focusing on a full chromosome-level assembly in future studies can potentially enhance the quality of the genome.

## Acknowledgments and Funding statement

The authors thank Anna Mácová and Roman HrdliČka for their invaluable assistance during the fieldwork. Additionally, we are grateful for the computational resources provided by the e-INFRA CZ project (ID: 90140), supported by the Ministry of Education, Youth and Sports of the Czech Republic. This research was made possible through the support of a grant from the Czech Science Agency (Grant No. GA19-04676S).

