## Supplemental Information for "Highly resolved genome assembly and comparative transcriptome profiling reveal genes related to developmental stages of tapeworm *Ligula intestinalis*"

Figure S1. Sampling localities in the Czech Republic and the life cycle of *L. intestinalis*.

Figure S2. Synteny and collinearity bar plot between the 10 longest scaffolds of *Ligula* genome and 200 and 203 scaffolds of *S. erinaceieuropaei* and *S proliferum*, respectively. Collinear gene blocks generated using Symap between genome scaffolds (>1 Mb) represent 18424, 8994, and 9324 genes for *L. intestinalis*, *S proliferum* and *S. erinaceieuropaei*, respectively.

Figure S3. The number of up and down regulated genes between life stages (Adult and Larva) and different parts of body (Apical, central, and posterior).

Table S1. Samples utilised for transcriptome data.

| Sample Id | Locality | Life Stage | Part of body | Number of reads | Fq Accession number |
| --- | --- | --- | --- | --- | --- |
| 9KLA | Klíčava | Plerocercoid | Anterior | 38,123,676 |  |
| 9KLC |  |  | Central | 27,183,954 |  |
| 9KLP |  |  | Posterior | 32,231,768 |  |
| 22KLA |  |  | Anterior | 32,479,215 |  |
| 22KLC |  |  | Central | 37,206,138 |  |
| 22KLP |  |  | Posterior | 30,784,170 |  |
| 35MA | Most |  | Anterior | 26,360,896 |  |
| 35MC |  |  | Central | 30,135,343 |  |
| 35MP |  |  | Posterior | 35,670,952 |  |
| 14M21A |  | Larva | Anterior | 30,693,070 |  |
| 14M21C |  |  | Central | 41,578,938 |  |
| 14M21P |  |  | Posterior | 29,247,701 |  |
| 13M21A |  |  | Anterior | 31,852,648 |  |
| 13M21C |  |  | Central | 28,431,333 |  |
| 13M21P |  |  | Posterior | 36,669,470 |  |
| 12M21A |  |  | Anterior | 29,620,438 |  |
| 12M21C |  |  | Central | 30,906,694 |  |
| 12M21P |  |  | Posterior | 35,670,952 |  |

Table S2. Statistics of sequencing data generated in this study.

|  | Nanopore | Novaseq | Omni-C |
| --- | --- | --- | --- |
| Total number of reads | 7,574,939 | 485,493,102 | 106,428,637 |
| Average read length | 5,699 | 150 | 150 |
| GC % | 45 | 44 | 44 |
| Assembly size,bp | 780,205,246 | 471,887,469 | 778,975,446 |
| Depth of Coverage (X) | 55 | 60 | 54 |
| Number of scaffolds | 137 | 81,578 | 61,514 |
| Longest Scaffold, bp | 2,329,270 | 143,614 | 1,726,504 |
| Scaffold N50 size, bp | 320,313 | 9,840 | 146,819 |

Table S3 Analysis of repeat content in the *Ligula* genome.

| Category | Number of elements | Length occupied | percentage of sequences |
| --- | --- | --- | --- |
| **Retroelements** | **450036** | **264202183 bp** | **34.07 %** |
| Penelope | 227627 | 86470978 bp | 11.15 **%** |
| LINEs | 397285 | 233051484 bp | 30.05 % |
| L2/CR1/Rex | 124069 | 112872190 bp | 14.55% |
| R1/LOA/Jockey | 475 | 511392 bp | 0.07 % |
| R2/R4/NeSL | 66 | 94388 bp | 0.01 % |
| RTE/Bov-B | 33424 | 23638860 bp | 3.05 % |
| LTR elements | 52751 | 31150699 bp | 4.02% |
| BEL/Pao | 5776 | 3525641 bp | 0.45 % |
| Gypsy/DIRS1 | 42744 | 27127907 bp | 3.50 % |
| **DNA transposons** | **51119** | **14165910 bp** | **1.83%** |
| hobo-Activator | 3898 | 1759147 bp | 0.23% |
| Tc1-IS630-Pogo | 14059 | 4797296 bp | 0.62 % |
| **Small RNA** | **15184** | **2946277 bp** | **0.38%** |
| **Low complexity** | **3913** | **233542 bp** | **0.03%** |
| **Simple repeat** | **53777** | **3099135 bp** | **0.40%** |
| **Unclassified** | **823742** | **189184304 bp** | **24.40%** |
| Total | 1397771 | 473831351 bp | 61.1% |
