## Supplementary figures and images for "Highly resolved genome assembly and comparative transcriptome profiling reveal genes related to developmental stages of tapeworm *Ligula intestinalis*"

### Figure S1

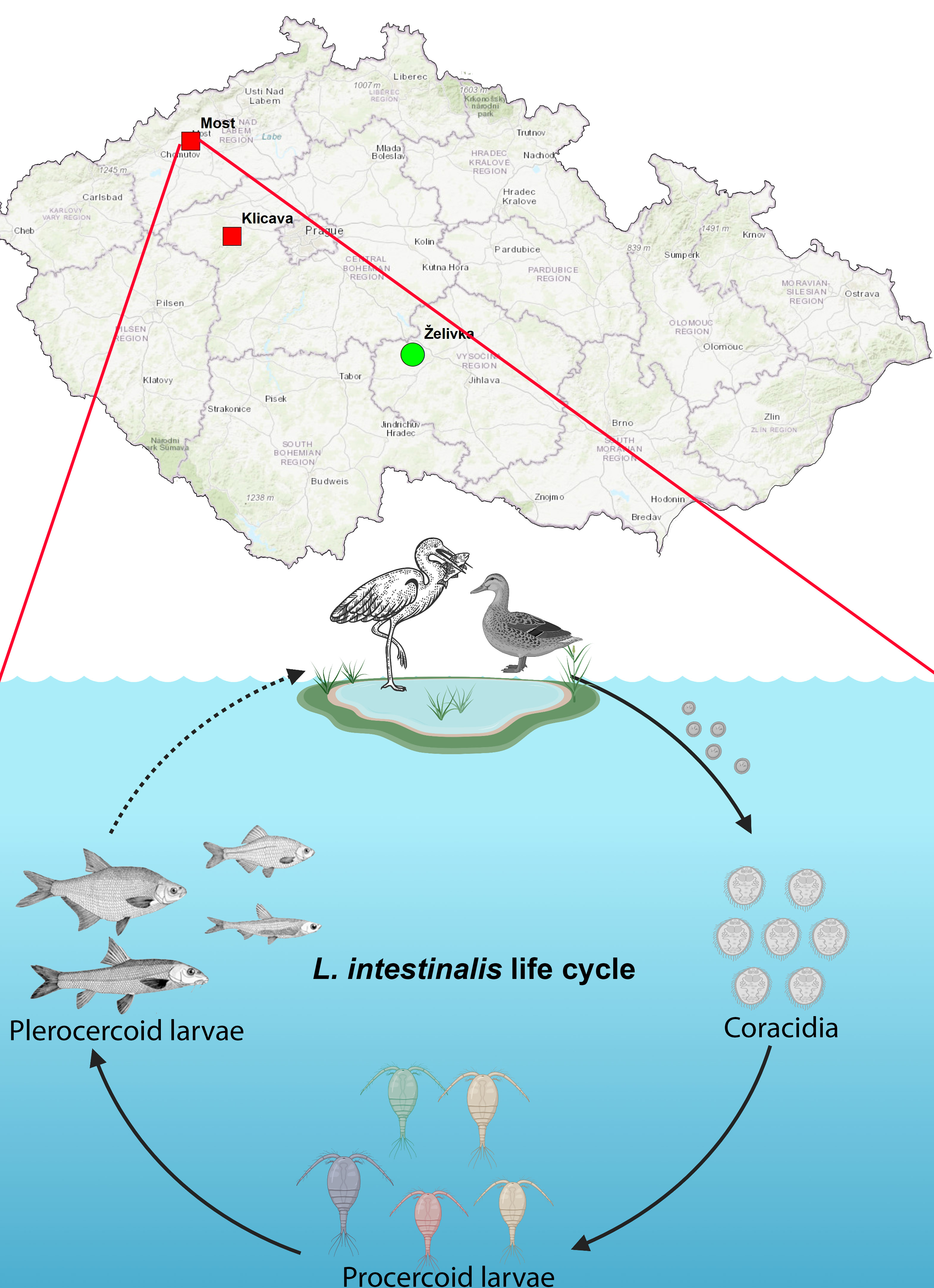

### Figure S2

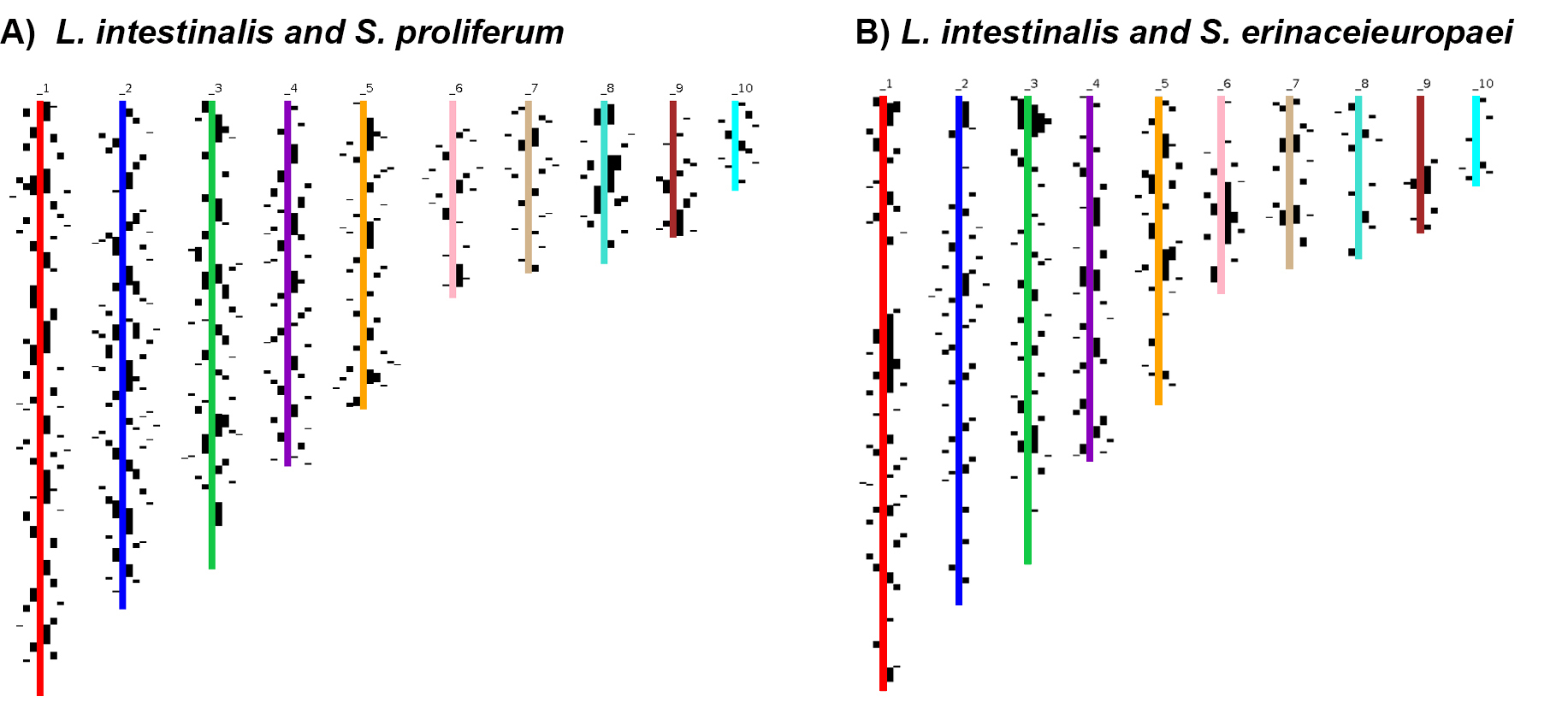

### Figure S3

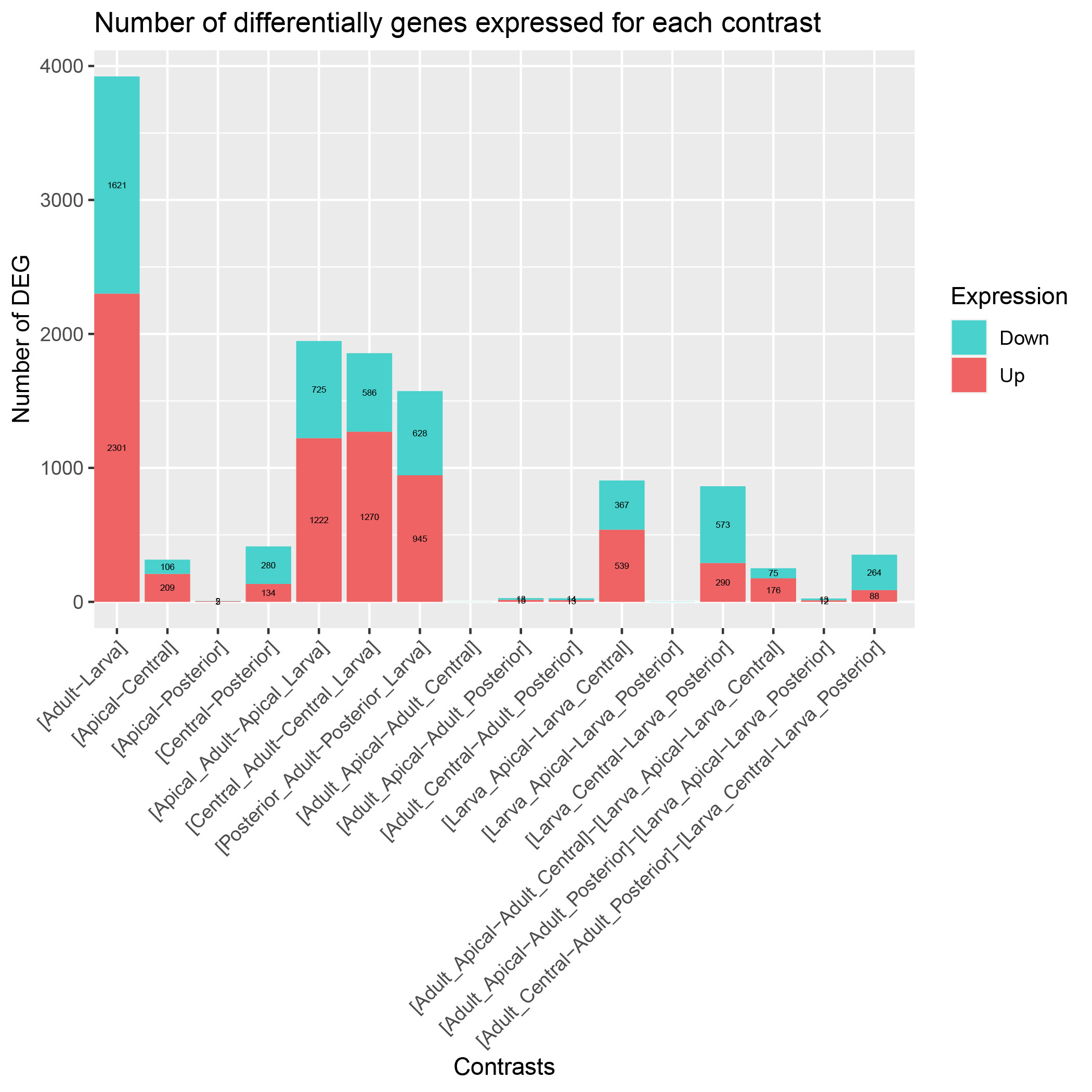
